# Easy MPE: Extraction of quality microplot images for UAV-based high-throughput field phenotyping

**DOI:** 10.1101/745752

**Authors:** Léa Tresch, Yue Mu, Atsushi Itoh, Akito Kaga, Kazunori Taguchi, Masayuki Hirafuji, Seishi Ninomiya, Wei Guo

## Abstract

Microplot extraction (MPE) is a necessary image-processing step in unmanned aerial vehicle (UAV)-based research on breeding fields. At present, it is manually using ArcGIS, QGIS or other GIS-based software, but achieving the desired accuracy is time-consuming. We therefore developed an intuitive, easy-to-use semi-automatic program for MPE called Easy MPE to enable researchers and others to access reliable plot data UAV images of whole fields under variable field conditions. The program uses four major steps: (1). Binary segmentation, (2). Microplot extraction, (3). Production of *.shp files to enable further file manipulation, and (4). Projection of individual microplots generated from the orthomosaic back onto the raw aerial UAV images to preserve the image quality. Crop rows were successfully identified in all trial fields. The performance of proposed method was evaluated by calculating the intersection-over-union (IOU) ratio between microplots determined manually and by Easy MPE: The average IOU (±SD) of all trials was 91% (±3).

## 1. Introduction

One of the major aims of investigations to improve agricultural machinery has been to reduce yield gaps [1]. The way we understand fields has been revolutionized by advances in yield monitoring technology, which have allowed field data to be measured with more and more precision. As a result, crop yield can now be measured at the microplot scale, and the same level of accuracy has been achieved by seeders and other kinds of agricultural machinery.

In recent years, great advances in sensors, aeronautics, and high-performance computing, in particular, have made high-throughput phenotyping more effective and accessible [2]. The characterization of quantitative traits such as yield and stress tolerance at fine-scale allows even complex phenotypes to be identified [3]. Eventually, specific varieties derived from a large number of different lines can be selected to reduce yield variation (e.g., [4–7]). Breeding efficiency now depends on scaling up the phenotype identification step, that is, on high-throughput phenotyping, as high-throughput genotyping is already providing genomic information quickly and cheaply [2].

Unmanned aerial vehicles (UAVs) have become one of the most popular information retrieval tools for high-throughput phenotyping, owing to the reduced costs of purchasing and deploying them, their easier control and operation, and their higher sensor compatibility. Proposed image analysis pipelines for extracting useful phenotypic information normally include several steps: 3D mapping by structure-from-motion (SfM) and multi-view stereo (MVS) techniques, generation of orthomosaic images (an image composed of several orthorectified aerial images stitched together) and digital surface models, extraction of microplots, and extraction of phenotypic traits. Among these steps, microplot extraction (MPE; i.e., cropping images to the size of individual subplots) is essential because it allows fields (or plots; that is to say cultured areas made of several microplots) to be examined at a level of detail corresponding to the level of accuracy of technologies such as the yield monitoring and seeder technologies mentioned above, ultimately providing better results [4, 8].

There are three main ways to perform MPE:

- Manually, usually by using geographic information system (GIS) software,
- Semi-automatically, which still requires some manual input,
- Automatically, which requires no manual input.

To the best of our knowledge, no fully automatic microplot extraction method has been published so far.

Several semi-automatic techniques have been developed to extract microplots from UAV images. A method proposed by Hearst [9] takes advantage of device interconnectivity by using the information on the geo-localization of crop rows from the seeder. The resulting orthomosaic thus contains spatial landmarks that allow it to be divided into microplots. However, this approach requires access to sophisticated technology and a high skill level. Most methods use image analysis tools, and they all require the user to define the maximum borders of one subplot. The whole field is then divided into equal-sized replicate subplots [10]. These methods require the plot to be very regular and consistently organized, which may not be the case, especially for breeding fields not planted by machine. More recently, Khan and Miklavcic [11] proposed a grid-based extraction method in which the plot grid is adapted to match the actual positions of plots. This technique allows for more field heterogeneity but it still requires the width/height, horizontal/vertical spacing, and orientation of a 2D vector grid cell to be specified interactively.

All of these techniques generally start with an orthomosaic image of the field rather than with raw images (i.e., unprocessed aerial images taken by the UAV, which are used to produce the orthomosaic). The consequent loss of image quality is an important consideration for high-throughput phenotyping.

Most MPE is still done manually, such as using shapefiles [12], the ArcGIS editor [13], the Fishnet function of Arcpy in ArcGIS [14], or the QGIS tool for plot extraction [15]. Manual extraction is not only time-consuming, depending on the size of the whole field, but also potentially less accurate because it is prone to human bias.

In this paper, we propose a semi-automated MPE solution based on image analysis. We tested two kinds of crops on the proposed method on six different field datasets. We chose sugar beet and soybean because of the importance of both crops are expected their unique growth patterns in each [22,23]. The growth pattern of the young plants fulfills the program requirements perfectly because they are generally planted in dense rows. Our ultimate goal was to elucidate the small differences in sugar beet and soybean growth in relation to microclimatic conditions in each field [24, 25]. Therefore, the accurate identification of micro plots allows the phenotypic traits of the crop in different locations of a field to be assessed, which could validate whether it was the correct program for high-throughput phenotyping of growth patterns or sizes correctly. First, as a pre-processing step, a binary segmented image is produced by one of two image segmentation methods: application of the Excess Green (ExG) vegetation index [16] followed by the Otsu threshold [17], or the EasyPCC program [18, 41]. The ExG index and the Otsu threshold have been shown to perform well at differentiating vegetation from non-plant elements (mostly bare soil) [19 – 21]. The EasyPCC program uses machine learning to associate pixels with either vegetation or background areas, based on the manual selection of example plants in the selected images; this program can produce a segmented image of a field when the ExG index and Otsu threshold method may not work, for example, because of light and color variation or other outdoor environmental variations on the image [19 – 21].

The pre-processing step is followed by MPE. To extract microplots, planted areas of the field are identified by summing the pixels classified as vegetation and then calculating the average. Microplot columns (90-degree rotation from crop rows) are identified by comparing vegetation pixel values with the average and erasing those below a threshold value. The same procedure is then applied to the column images to identify crop rows within each microplot column. The initial orthomosaic is then cropped to the borders identified by the program and a shapefile is generated for each microplot to facilitate further image manipulation in GIS software or custom programs. Microplot intersection points are also saved in an Excel spreadsheet file.

Finally, if raw unprocessed aerial images and SfM files (containing information on internal and external camera parameters) are available, the program can directly extract the identified microplots from the raw images so that the microplots will contain the maximum level of detail.

## 2. Materials and Methods

### 2.1. Experimental Fields and Image Acquisition

Six field datasets were used as experimental units (Table 1, Table S1). The plants were sown in precise plot configurations, and all trial fields were managed according to the ordinary local management practices.

**Table 1:**
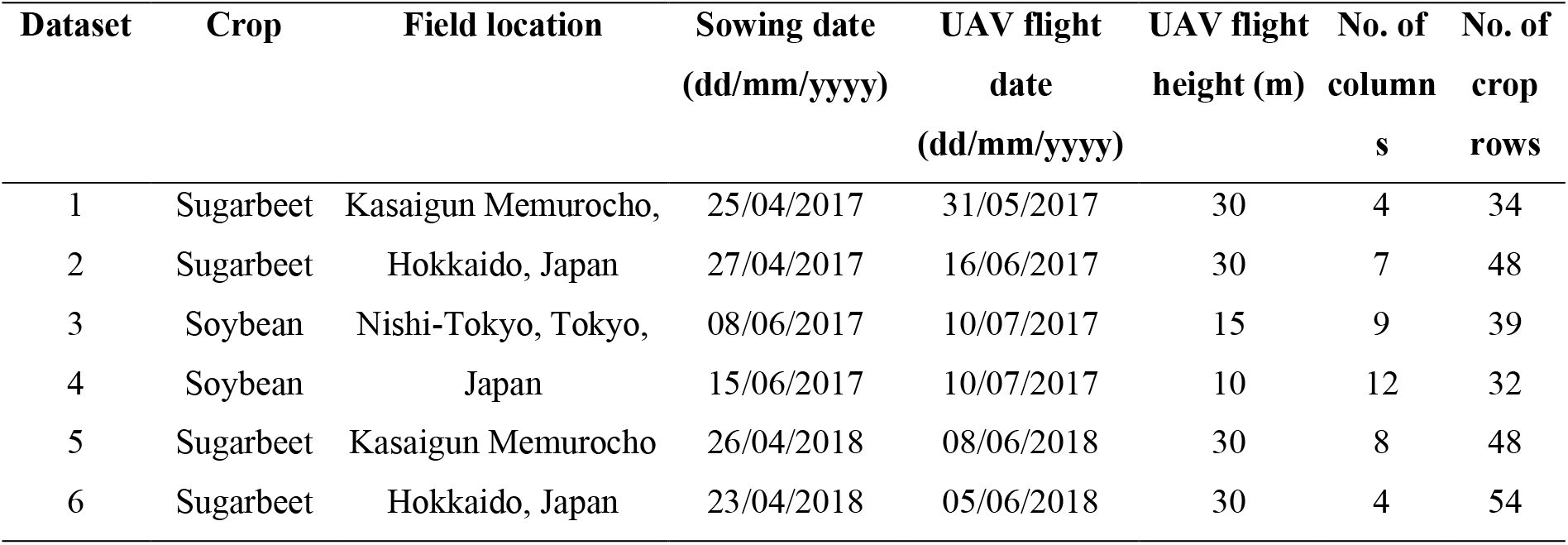
Trial field and image acquisition information.

| **Dataset** | **Crop** | **Field location** | **Sowing date**<br>**(dd/mm/yyyy)** | **UAV flight**<br>**date**<br>**(dd/mm/yyyy)** | **UAV flight**<br>**height (m)** | **No. of**<br>**column**<br>**s** | **No. of**<br>**crop**<br>**rows** |
| --- | --- | --- | --- | --- | --- | --- | --- |
| 1 | Sugarbeet | Kasaigun Memurocho, | 25/04/2017 | 31/05/2017 | 30 | 4 | 34 |
| 2 | Sugarbeet | Hokkaido, Japan | 27/04/2017 | 16/06/2017 | 30 | 7 | 48 |
| 3 | Soybean | Nishi-Tokyo, Tokyo, | 08/06/2017 | 10/07/2017 | 15 | 9 | 39 |
| 4 | Soybean | Japan | 15/06/2017 | 10/07/2017 | 10 | 12 | 32 |
| 5 | Sugarbeet | Kasaigun Memurocho | 26/04/2018 | 08/06/2018 | 30 | 8 | 48 |
| 6 | Sugarbeet | Hokkaido, Japan | 23/04/2018 | 05/06/2018 | 30 | 4 | 54 |

Before image acquisition, the waypoint routing of the UAV was specified by using a self-developed software tool that automatically creates data formatted for import into Litchi for DJI drones (VC Technology, UK). This readable waypoint routing data allowed highly accurate repetition of flight missions so that all images had the same spatial resolution. Orthomosaics of the georeferenced raw images were then produced in Pix4Dmapper Pro software (Pix4D, Lausanne, Switzerland; Table S1).

UAVs overflew each field several times during the growing period of each crop. The flight dates in Table 1 are dates on which images that suited the program requirements were obtained (i.e., with uniform crop rows that did not touch each other and low content of weeds).

### 2.2. Easy Microplot Extraction

The Easy MPE code is written in the Python language (Python Software Foundation, Python Language Reference, v. 3.6.8 [26]) with the use of the following additional software packages: OpenCV v. 3.4.3 [27]; pyQt5 v. 5.9.2 [28]; NumPy v. 2.6.8 [29], Scikit-Image v. 0.14.1 [30]; Scikit-Learn v. 0.20.1 [31]; pyshp package v. 2.0.1 [32]; rasterio v. 1.0.13 [33], rasterstats v. 0.13.0 [34]; and Fiona v. 1.8.4 [35].

The main aim of the program is to identify columns, and crop rows within each column, on an orthomosaic image of a field so that microplot areas can be extracted. Additional outputs that the user may need for further analysis of the data are also provided. The program comprises four processing steps: binary segmentation (pre-processing); microplot identification and extraction; production of shapefile and additionally production of microplots extracted from raw individual images by reverse calculation.

The code was run on a laptop computer (Intel Core i7-8550U CPU @ 1.80 GHz, 16 GB RAM, Windows 10 Home 64-bit operating system) connected to an external graphics processing unit (NVIDIA, GeForce GTX 1080Ti).

Figure 1 shows the four major steps of the Easy MPE program, and the binary segmentation and microplot extraction details are described in smaller steps A–I. Orange arrows indicate automated steps. In step A to B, however, the region of interest in the orthomosaic must be manually selected. Note that for simplicity and clarity, some steps have been omitted.

**Figure 1:**
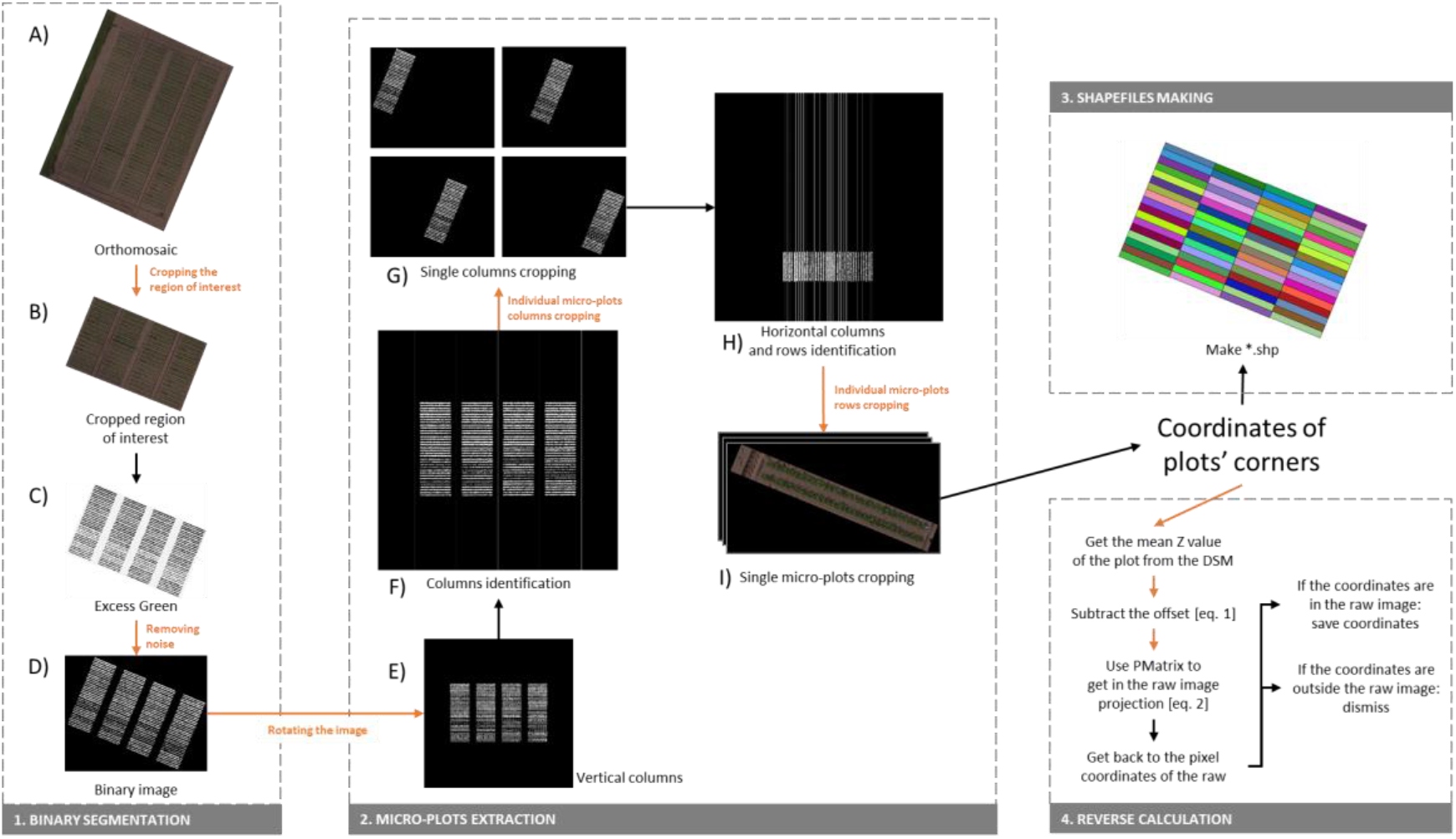
Global pipeline of the Easy MPE program, demonstrated using dataset 3.

#### 1. Binary segmentation

The ExG vegetation index (step C) and the Otsu threshold (step D) are applied to each pixel of an orthomosaic image to produce a segmented binary image (Figure 1). Excess Green is calculated as:

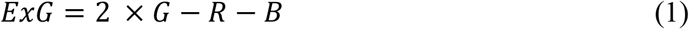

where *R*, *G*, and *B* are the normalized red, green, and blue channel values of a pixel.

In the case of a very bright or very dark environment, where the Otsu threshold does not perform well [19] [20] [21], EasyPCC is used [18].

#### 2. Microplot extraction

MPE is the core step of the program. In this step, binary images of diverse field layouts are segmented (partitioned) into columns and crop rows.

First, the image is rotated (step E) using the coordinates of the bounding box drawn around the binary image. In step A, it is important to draw the boundaries of the region of interest so that no object that might be segmented lies between the boundaries and the crop; otherwise, the image will not be correctly rotated.

Second, columns are extracted by identifying local maxima along the *x*-axis of the image, based on the sum of the white pixels distributed in the columns. To make the columns even, the binary image is slightly manipulated with an erosion followed by a horizontal dilatation. White pixels along the *x*-axis are first summed and their average is calculated. Then all values that are less than ⅓ of the average are erased. The erased pixels represent “transition parts” at either end of the crop rows as well as small intercolumn weeds. Columns are then identified (step F) by scanning along the *x*-axis: a transition from black to white pixels marks the left edge of a column, and the subsequent transition from white to black pixels marks its right edge.

Third, on the orthomosaic binary image, crop rows are identified within each column (step H). Because crop rows are more homogeneous along the *x*-axis than the columns identified in step F, the intra-row variation in the sum of the white pixels is expected to be small; moreover, inter-row weeds are expected to be frequent. Thus, values of less than ½ of the average sum are erased in step H. Then crop rows are identified by scanning along the y-axis and identifying transitions in the segmented image from black to white or from white to black pixels.

According to the number of crop rows and the number of columns in each field input into the program, microplots are cropped and saved (step G). Their coordinates are calculated in the orthomosaic coordinate system, or if the orthomosaic does not include coordinates, from the positions of the pixels on the image.

#### 3. Shapefile production and Reverse calculation

The program can output shapefiles of the microplots.

Shapefiles are produced by using the coordinates calculated during the MPE step, with the same characteristics as the orthomosaic image.

Reverse calculation is the process of projecting individual microplots determined from the orthomosaic image back onto the corresponding area of the raw images. It aims to preserve the raw image resolution, instead of the lower quality of the orthomosaic image (Figure 2), for estimation of phenotypic traits such as ground coverage [8].

**Figure 2:**
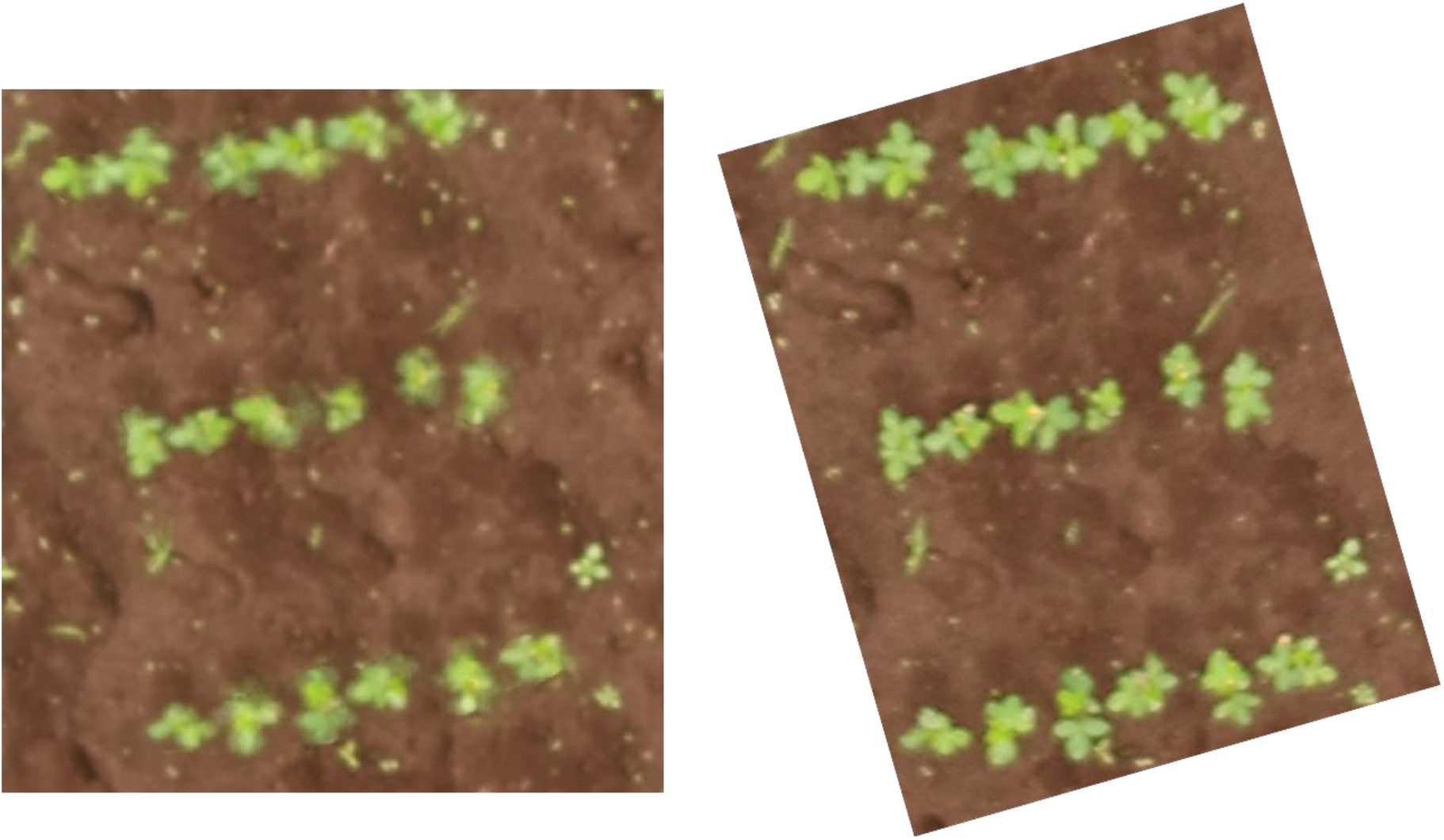
Quality diminution in an orthomosaic from dataset 3 (left) compared to the orthomosaic (right)

For the field trials, the following Pix4Dmapper Pro [36] outputs (Table 1) were used: P-Matrix (contains information on the internal and external camera parameters), offset (the difference between the local coordinate system of the digital surface model (DSM] and the output coordinate system), the DSM, and the raw image data used to produce an orthomosaic of the field. Equations (2) – (5) are used to determine the coordinates of each microplot in the raw images:

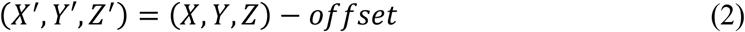

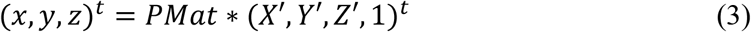

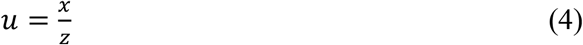

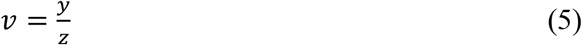

(*X, Y, Z*) are the 3D coordinates of the microplot: the values of *X* and *Y* are obtained during the microplot extraction step, and *Z* is the average height of the microplot in the DSM. (*X’, Y’, Z’*) are the corrected 3D coordinates of the microplots after they have been fit to the P-Matrix (PMat) coordinates. (*x, y, z*) are intermediate 3D coordinates in the camera’s coordinate system that are used to convert 3D points into 2D points. (*u, v*) are the 2D pixel coordinates of the microplot in the raw image.

Easy MPE outputs a “*.csv” file with 11 columns (column number, crop row number, raw image name, and the four (*u, v*) corner coordinates of the microplot in the raw image). This output was chosen to exclude unwanted data that would unnecessarily increase computation times while including the information necessary for the user to be able to easily use the data.

Finally, computational times were determined by measuring the processing time required for each major step, because manual inputs are required in between steps. Processing times were determined with the time module implemented in Python five times for each field.

### 2.2. Parameters Input into the MPE Program

The input parameters of the MPE program are field orthomosaic, type of image (binary or RGB), noise, number of columns, number of crop rows per column, and global orientation of the columns in the orthomosaic (horizontal or vertical). The noise parameter is used after the segmentation. Objects smaller than the input value are removed in order to output a homogeneous image without signals labeled as noise by the user.

For each dataset, the inputs were either self-evident (type of image, number of columns, number of crop rows per column, global orientation of the columns) or determined from the field conditions and image resolution (image input, noise). The targeted area in the orthomosaic (i.e., the field) had to be manually delimited so that the program would be applied to the desired area.

Datasets 5 and 6 were binary images and datasets 1 to 4 were RGB images. One image, dataset 4, had to be resized because it was too large for our computer to handle.

Because the targeted area was manually delimited by the user, the program outputs could change slightly between trials.

Input details are available in **Table S2**.

### 2.3. Manually Produced Reference Microplots

To evaluate the performance of Easy MPE, we produced reference microplots manually and used them as ground-truth data. The desired program output consists of microplots, each having the number of columns and crop rows specified by the user, so that the user’s experimental design can be fit to the field or so that local field information can be determined more precisely.

Reference microplots were delimited by hand on orthomosaic images, as precisely as possible until the level of accuracy was judged by the user to be sufficient. We aimed at minimizing the exclusion of the target-MPE plant pixels and minimizing the inclusion of adjacent rows or columns. Shapefiles for the reference images were produced by using the grid tool in QGIS Las Palmas v. 2.18.24 software [37]. The free, open-source QGIS was used because it is intuitive and accessible to all users.

Although the manually produced reference microplots may include bias introduced by the user, their use does not compromise the study’s aim, which was to compare program-delimited microplots with manually delimited ones.

### 2.4. Performance Evaluation: Intersection Over Union

The manually determined microplots were automatically compared with the program-generated microplots by using the saved coordinates in their shapefiles and a Python program. This program uses pyshp v. 2.0.1 [32] to retrieve the *.shp coordinates from the shapefiles and shapely v. 1.6.4 [38] to compare the microplot areas.

We used the intersection-over-union (IOU) performance criterion [39] to evaluate the similarity between the predicted area and the ground-truth area of each microplot. IOU is a standard performance measure used to evaluate object category segmentation performance (Figure 3).

**Figure 3:**
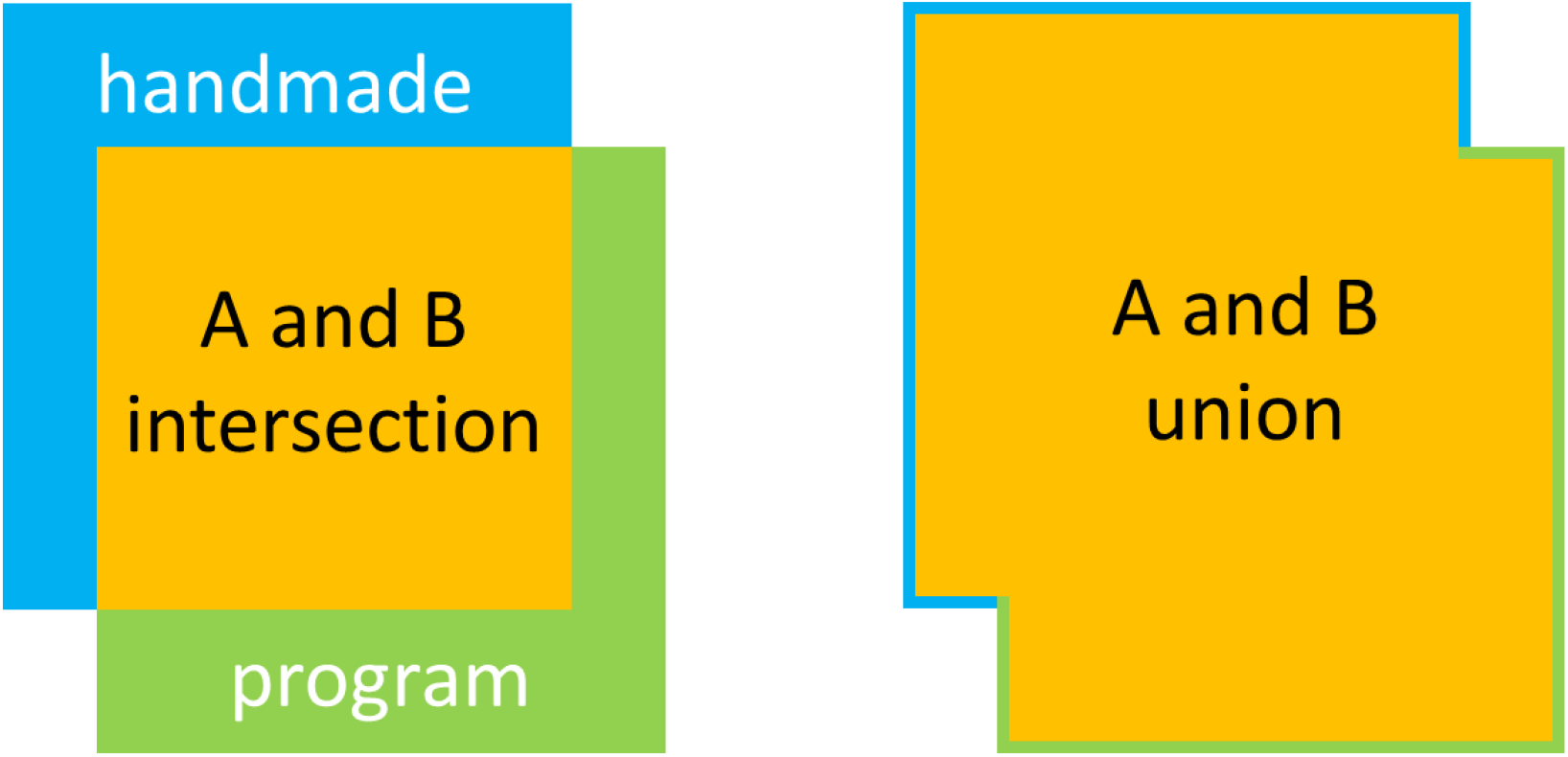
Visual representation of the intersection (yellow area on the left) and union (yellow area on the right) areas of manually (blue) and program-determined (green) areas.

IOU is calculated as (6):

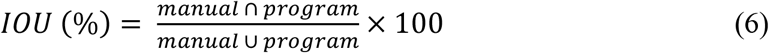

The output is a *.csv file with six columns: plot identification number, the program *.shp area, the manually determined *.shp area, the intersection area, the union area, and the IOU.

### 2.5. Statistical Analysis

The population standard deviation is calculated for the IOU as (7):

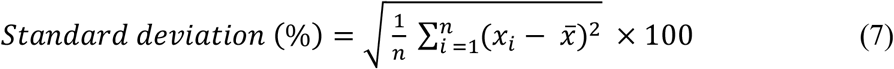

where *n* is the number of microplots in the targeted field, *x_*i*_* is the IOU ratio of microplot *i*, and *x̅* is the mean IOU of the targeted field.

Precision *P* and recall *R* are defined by equations (8) and (9), respectively:

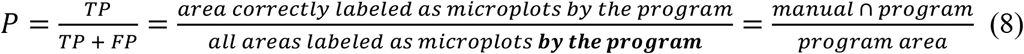

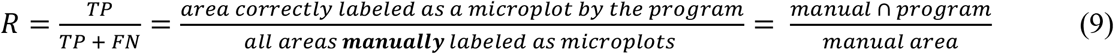

TP indicates a true positive: what has been correctly considered by the program to be part of a microplot; thus, TP = manual area ∩ program area (yellow area on the left side of Fig. 3). FP indicates a false positive: what has been incorrectly been considered by the program to be part of a microplot; thus FP = program area – manual ∩ program area (green area on the left side of Fig. 3). FN means a false negative: areas in the manually determined shapefiles that were not considered to be part of microplots in the program shapefiles. Thus, FN = manual area – manual ∩ program area (blue area on the left side of Fig. 3).

Thus, *P* gives the percentage of the program-determined microplot area that has been correctly identified by the program, and *R* is the percentage of the manually determined microplot area that has been correctly identified by the program. If the program-determined (manually determined) area is wholly included in the manually determined (program-determined) area, then *P* (*R*) will be equal to 100%.

We also calculated the standard deviations of *P* and *R*.

## 3. Comparison results

**Figure 4:**
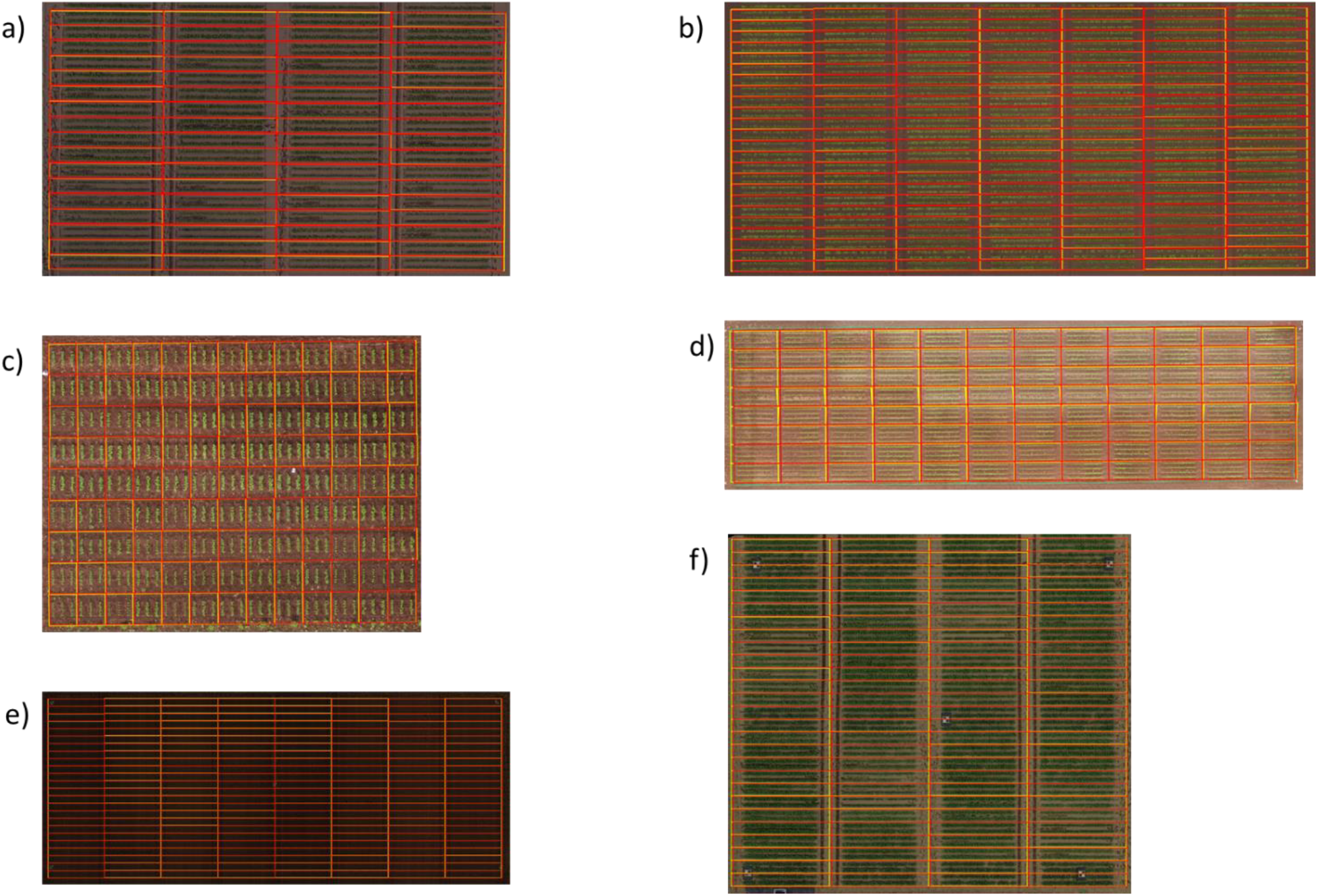
Comparisons of manually determined (yellow lines) and program-determined (red lines) microplot boundaries: (a – f) Datasets 1–6.

**Table 2:** Intersection-over-union results.

| Trial fields | Average IOU (%) | SD of IOU (%) | Average precision (%) | SD of precision (%) | Average recall (%) | SD of recall (%) |
| --- | --- | --- | --- | --- | --- | --- |
| Dataset 1 | 93.0 | 3.0 | 95.0 | 2.20 | 98.0 | 2.08 |
| Dataset 2 | 86.0 | 5.0 | 95.0 | 2.22 | 91.0 | 5.64 |
| Dataset 3 | 92.0 | 3.0 | 96.0 | 1.86 | 96.0 | 2.06 |
| Dataset 4 | 88.0 | 3.0 | 94.0 | 2.09 | 94.0 | 1.62 |
| Dataset 5 | 93.0 | 4.0 | 97.0 | 2.18 | 96.0 | 2.04 |
| Dataset 6 | 92.0 | 3.0 | 96.0 | 1.72 | 96.0 | 1.87 |
| **Average** | **90.7** | **3.5** | **95.5** | **2.05** | **95.2** | **2.55** |

The program successfully identified microplots in all trial fields (Fig. 4).

The mean IOU among the datasets was 91% (±%3), indicating a 91% overlap between program-determined and manually determined microplot areas. Moreover, among these fields, an IOU of < 80% was obtained only for dataset 2, which comprised 20 individual microplots (the lowest IOU being 71%).

Mean precision and recall were 95% (±2%) and 95% (±2%), respectively. These results indicate that neither manually determined nor program-determined microplots showed a tendency to be wholly included in the other. Therefore, the IOU can be understood to indicate a shifting of microplot boundaries between them.

### 3.2. Computation Time

Computation time depends, of course, on the computer used to run the program.

The computational times (Table 3) varied among the trial images, but some global trends can be observed by comparing the computational times in Table 3 with dataset information provided in Table S1 and S2. The program was slower overall when dealing with larger images, which is to be expected because the program involves image manipulation. The computational time required for binary segmentation depended mostly on the type of input (binary or RGB), whereas the time required for microplot extraction and reverse calculation depended mainly on the number of microplots in the trial field, because this number determines the number of images that must be manipulated in each of these steps.

**Table 3:** Average computational times of Easy MPE per major step for each dataset.

| Trial field | Binary segmentation<br>(s) | Microplot extraction<br>(s) | Reverse calculation (s) | Total time (s) |
| --- | --- | --- | --- | --- |
| Dataset 1 | 8.537 | 97.925 | 22.75 | 129.212 |
| Dataset 2 | 15.491 | 551.794 | 80.76 | 648.045 |
| Dataset 3 | 16.609 | 403.081 | 60.7 | 480.39 |
| Dataset 4 | 22.972 | 670.2 | * | * |
| Dataset 5 | 2.631 | 191.369 | 65.419 | 259.419 |
| Dataset 6 | 1.506 | 68.485 | 20.272 | 90.263 |

\* Reverse calculation could not be performed on dataset 4 due to a lack of the required inputs.

## 4. Discussion

The microplot results obtained by Easy MPE were similar to those obtained manually. However, manual identification means only that the accuracy is controlled and judged to be acceptable. Thus, the relationship between the IOU and “absolute accuracy” depends on the precision with which the reference plots are defined. The study results confirm that the accuracy of the program is similar to that obtained by manual identification of microplots, but a direct comparison of the results obtained by the two methods suggests that the program-determined microplots are more precise and more regular than those determined manually (Fig. 4).

In addition, Easy MPE places a few conditions on the initial image. The field must be fairly homogeneous and crop rows should not touch each other, or not much. These conditions were met by both the soybean and sugarbeet fields in this study, but it is necessary to test other types of fields as well. Continuous field observations (including multiple drone flights) are likely to be required to get the usable image. Image pretreatment by removing weed pixels, either manually or automatically, would help Easy MPE to get good results.

The only manual input to the program is plot delimitation by the user at the very beginning. It is essential that no segmented objects other than the field be included in the region of interest. It might be possible to automate this step by providing GPS coordinates from the seeder or by using an already cropped orthomosaic image, but either method would diminish the freedom of the user to apply the program to an area smaller than the whole field.

Also please note that application of the program is limited by the available computer, as shown by the example of dataset 4.

The MPE method used by Easy MPE is different from previously published methods. Easy MPE uses image parameters and information on the field geometry to adapt itself to the field, whereas in the method proposed by Khan and Miklavcic [12], a cellular grid is laid over the image so that each individual rectangular cell is optimally aligned. Online software (e.g., [11]) also uses a grid and requires the user to indicate the positions of several microplots. Easy MPE asks for a working zone delimitation and no other image manipulation, increasing its user-friendliness. This method has been built as a piece of a suite of open-source programs for high-throughput phenotyping, along with Easy PCC [18] and future additions.

Other published methods do not include the reverse calculation step, which allows Easy MPE to provide images of the same quality as the raw images. The Easy MPE reverse calculation procedure is coded for Pix4D outputs, but outputs of free SfM software outputs could be used as well. Three files are required [40]:

- The P-Matrix file, which contains information about the internal and external camera parameters, allowing the 3D coordinates to be converted into 2D coordinates.
- A DSM file, which is the usual output of SfM software.
- An Offset file, which contains the offsets between local coordinate system points used in the creation of some of the output and the output coordinate system.

Note that the code would need to be adapted to be able to extract the needed data from free SfM software outputs, but the adapted Easy MPE code would then be entirely free.

Other possible improvements include the implementation of a different vegetation index such as NDVI, SAVI, or GNDVI for preprocessing segmentation, and the linking of EasyPCC to Easy MPE. Verification methods could also be added as an option; for example, the Hough transform or the average distance between crop rows within microplots could be used to verify that the field components are correctly identified. These methods could be used only with geometric fields that are very constant in the seeds repartition; they would thus narrow the applicability of the code.

Finally, the code needs to be further tested and improved by applying it to other important crops that are not as densely planted as soybean and sugarbeet, such as maize or wheat. Fields with crop-residues have not been tested either in this demonstration. The impact of a poor quality orthomosaic has not been investigated and should be measured in order to give the plant phenotyping community good insights about the possible uses of EasyMPE.

Overall:

- Easy MPE is recommended for its user-friendliness and simplicity; however, many points still have to be tested and approved, which leaves room for improvement. The micro-plots are delimited in an unbiased way, i.e. without human influence. It is part of an open-source suite of programs designed for the plant phenotyping community, include a segmentation step if needed and provides the first automation of the reverse calculation process.
- Grid-based programs are efficient as demonstrated by many publications. Improvement has been made recently, as in [11], and gives quite robust tools for MPE identification. It automatically requires for crops to be rectangular-shaped and adding a grid can be impacted by human perception, adding possible errors. It does not provide any additional services.
- In [9], the process can be fully automatic if the process and GPS localization are extremely precise (RTK recommended) and the field can access a high level of technology and informatics competencies.

## Acknowledgments

The authors thank Dr. Shunji Kurokawa, Division of Crop Production Systems, Central Region Agricultural Research Center, NARO, Japan; technicians of the Institute for Sustainable Agro-ecosystem Services, The University of Tokyo; and members of the Memuro Upland Farming Research Station, Hokkaido Agricultural Research Center, NARO for field management support.

## Author contributions

LT developed the algorithm and Python code with input from WG; YM conducted the reverse calculations; AI developed the executable Windows program; WG, AK, and KT conceived, designed, and coordinated the field experiments; WG, MH, and SN supervised the entire study; LT wrote the paper with input from all authors. All authors read and approved the final manuscript.

## Data Availability

Submission of a manuscript to Plant Phenomics implies that the data is freely available upon request or has deposited to a open database, like NCBI. If data are in an archive, include the accession number or a placeholder for it. Also include any materials that must be obtained through an MTA.

https://github.com/oceam/EasyMPE

## Supplementary Materials

**Table S1:**
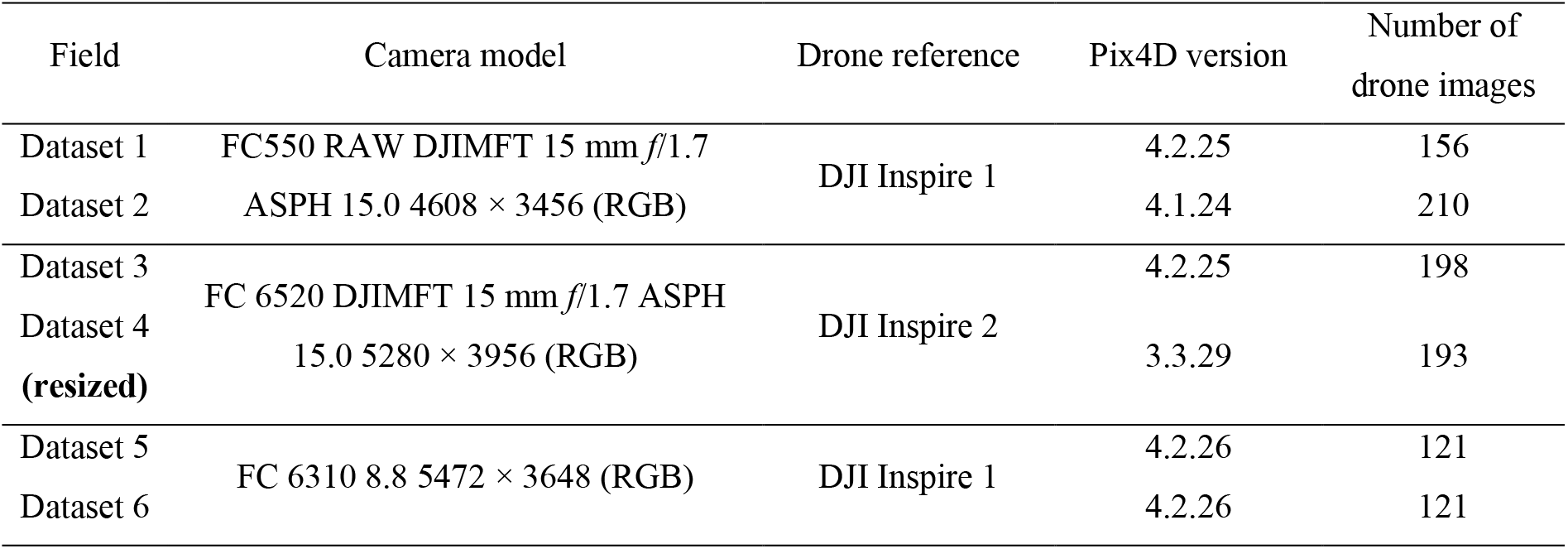
Details about image acquisition of the trial fields.

**Table S2:** Easy MPE program inputs for each dataset.

| Trial field | Type of image | Date of chosen image<br>(dd/mm/yyyy) | Noise (px) | Number of columns per microplot | Number of crop rows per microplot column | Column orientation |
| --- | --- | --- | --- | --- | --- | --- |
| Dataset 1 | RGB | 31/05/2017 | 200 | 1 | 2 | Vertical |
| Dataset 2 | RGB | 16/06/2017 | 100 | 1 | 2 | Vertical |
| Dataset 3 | RGB | 10/07/2017 | 1000 | 1 | 3 | Vertical |
| Dataset 4 | RGB | 10/07/2017 | 200 | 1 | 4 | Horizontal |
| Dataset 5 | Binary | 07/06/2018 | 700 | 1 | 2 | Vertical |
| Dataset 6 | Binary | 05/06/2018 | 500 | 1 | 2 | Vertical |

